# A site-wise reliability analysis of the ABCD diffusion fractional anisotropy and cortical thickness: impact of scanner platforms

**DOI:** 10.1101/2024.02.21.581460

**Authors:** Yezhi Pan, L. Elliot Hong, Ashley Acheson, Paul M. Thompson, Neda Jahanshad, Alyssa H. Zhu, Jiaao Yu, Chixiang Chen, Tianzhou Ma, Ho-Ling Liu, Jelle Veraart, Els Fieremans, Nicole R. Karcher, Peter Kochunov, Shuo Chen

## Abstract

The Adolescent Brain and Cognitive Development (ABCD) project is the largest study of adolescent brain development. ABCD longitudinally tracks 11,868 participants aged 9-10 years from 21 sites using standardized protocols for multi-site MRI data collection and analysis. While the multi-site and multi-scanner study design enhances the robustness and generalizability of analysis results, it may also introduce non-biological variances including scanner-related variations, subject motion, and deviations from protocols. ABCD imaging data were collected biennially within a period of ongoing maturation in cortical thickness and integrity of cerebral white matter. These changes can bias the classical test-retest methodologies, such as intraclass correlation coefficients (ICC). We developed a site-wise adaptive ICC (AICC) to evaluate the reliability of imaging-derived phenotypes while accounting for ongoing brain development. AICC iteratively estimates the population-level age-related brain development trajectory using a weighted mixed model and updates age-corrected site-wise reliability until convergence. We evaluated the test-retest reliability of regional fractional anisotropy (FA) measures from diffusion tensor imaging and cortical thickness (CT) from structural MRI data for each site. The mean AICC for 20 FA tracts across sites was 0.61±0.19, lower than the mean AICC for CT in 34 regions across sites, 0.76±0.12. Remarkably, sites using Siemens scanners consistently showed significantly higher AICC values compared to those using GE/Philips scanners for both FA (AICC=0.71±0.12 vs 0.46±0.17, p<0.001) and CT (AICC=0.80±0.10 vs 0.69±0.11, p<0.001). These findings demonstrate site-and-scanner related variations in data quality and underscore the necessity for meticulous data curation in subsequent association analyses.

## 1. Introduction

Adolescence is a crucial period for brain development that is associated with myelination of cerebral white matter tracts and pruning of cortical grey matter that support development of higher cognitive functions (Gogtay, Giedd et al. 2004, Gogtay and Thompson 2009, Bartzokis, Lu et al. 2010, Kochunov, Williamson et al. 2012, Kochunov, Thompson et al. 2015). Adolescence is also associated with the onset of symptoms for severe mental illnesses (SMI), such as schizophrenia, bipolar, or major depressive disorders, substance use and others (Rapoport, Addington et al. 2005, Casey, Nigg et al. 2007, Kalia 2008). Adolescent Brain Cognitive Development (ABCD) is the largest longitudinal study of brain development and child health consisting of N=11,868 participants aged 9-10 years at baseline, ascertained at 21 sites across the US (Karcher and Barch 2021). The ABCD collection neuroimaging approaches were developed to collect data for quantitative longitudinal analysis of multi-site diffusion, structural and functional maturational changes (Casey, Cannonier et al. 2018, Hagler, Hatton et al. 2019) . This included standardized imaging protocols and preprocessing pipelines designed for multi-site homogenization and phenotype extraction (Casey, Cannonier et al. 2018, Hagler, Hatton et al. 2019). However, non-biological variations were reported in ABCD data due to differences in scanners, deviations from the protocol, imaging artifacts and participant motion (Nielson, Pereira et al. 2018). Manual quality control of the ABCD T1-weighted images suggested that up to 50% of the scans were affected with non-biological variance (Elyounssi, Kunitoki et al. 2023). Here, we performed quality assessment of longitudinal regional measurements of fractional anisotropy (FA) of water diffusion extracted for major white matter tracts using the ABCD recommended pipeline. We specifically evaluated site-related differences in longitudinal fidelity of the FA values, including the differences in data collected using 3T scanners manufactured by Siemens, Philips, and General Electrics (GE). We developed an adaptive intraclass correlation coefficient (AICC) measure to evaluate the test-retest reliability of imaging-derived phenotypes while accounting for brain developmental trends in the population.

We focused on evaluating the impact of non-biological variance on the longitudinal DTI-FA measurements of cerebral white matter (WM). FA is important because it is a sensitive biomarker for non-invasive study of WM development (Basser 1994, Ulug, Barker et al. 1995, Conturo, McKinstry et al. 1996, Pierpaoli and Basser 1996). FA values are sensitive to many parameters (Beaulieu 2002) but the longitudinal changes in regional FA values during normal maturation are thought to be predominantly due to changes in myelination (Song, Sun et al. 2003, Song, Yoshino et al. 2005, Budde, Kim et al. 2007, Madler, Drabycz et al. 2008, Ryan, Sherman et al. 2017, Ryan, Sherman et al. 2018). The regional changes in cerebral FA values were used to replicate classical findings by Flechsig who demonstrated that continued myelination of WM during adolescence and early adulthood is the basis for development of higher cognitive function (Flechsig 1901, Kochunov, Williamson et al. 2012). Herein, we evaluated the ability to detect longitudinal changes in FA, compared with longitudinal changes in cortical gray matter thickness and speculated on potential causes of non-biological variances.

There is no single established metric for evaluating test-retest reproducibility of neuroimaging measurements in longitudinal developmental studies. Many studies used intraclass correlation coefficients (ICC) to demonstrate reproducibility for metrics such as FA, cortical thickness, and other measurements (Shrout and Fleiss 1979, Wijtenburg, Gaston et al. 2013, Zuo and Xing 2014, Acheson, Wijtenburg et al. 2017, Xue, Yuan et al. 2021). However, ICC and other approaches may not be applicable to the ABCD data because the neuroimaging measures are collected biennially at the time when participants are undergoing rapid development (Kochunov, Glahn et al. 2011, Kochunov, Glahn et al. 2011, Kochunov, Williamson et al. 2012). ICC is performed with the assumption of repeated measures performed under similar conditions, and the dynamic alterations in the developing adolescent brain will bias the reliability measures (Barnea-Goraly, Menon et al. 2005, Barnhart, Haber et al. 2007, Casey, Jones et al. 2008, Konrad, Firk et al. 2013). In the present work, we describe an AICC measure to evaluate the reliability of imaging-derived phenotypes acquired from the ABCD study while accounting for the normative age-related changes. We first estimate developmental trajectory using the complete dataset, for it is more robust and accurate than site-wise estimation. The iterative AICC estimation process also factors site-wise data reliability into the calculation of age effects. The resulting site-wise AICC can be integrated into subsequent statistical analyses to reduce bias and enhance inference efficiency. A simulation study was also conducted to assess the accuracy and robustness of our method.

## 2. Materials and Methods

### 2.1 Study samples

This study used baseline, two-year, and four-year follow up data from the NIMH Data Archive ABCD Curated Data Release 5.0 (https://abcdstudy.org/). The cohort and study protocols can be found in Garavan et al. (Garavan, Bartsch et al. 2018). Overall, the ABCD release 5.0 included early longitudinal data on 11,868 demographically diverse subjects, including neuroimaging data and other phenotypic data. For inclusion in the analyses, subjects were required to have both imaging data and relevant imaging acquisition information available. Additionally, subjects meeting any of the following exclusion criteria were excluded for the evaluation of site-wise data reliability: 1) attendance at different study sites during follow-up visits; 2) absence of longitudinal information.

### 2.2 ABCD image acquisition

In the ABCD study, imaging data were acquired using Siemens (Prisma VE11B-C), Philips (Achieva dStream, Ingenia), and GE (MR750, DV25-26) 3-Tesla MRI scanners. Siemens scanners were equipped with either 32 or 64 channel head coils. Philips scanners used 32 channel head coil. GE protocol required the use of Nova Medical 32 channel coil. The scanner-specific sequences and sequence parameters were standardized across different scanner platform with exception of minor discrepancies due to the hardware and software constraints (Casey, Cannonier et al. 2018, Hagler, Hatton et al. 2019).

#### 2.2.1 Siemens scanners

For the T1-weighted session, a matrix of 256×256, 176 slices, 1.0mm isotropic resolution, a repetition time (TR) of 2500ms, echo time (TE) of 2.88ms, and a field of view (FOV) of 256×256 were used. T2-weighted employed the same matrix, slices, and resolution, with a TR of 3200ms and a TE of 565ms. The dMRI session had a matrix of 140×140, 81 slices, 1.7mm isotropic resolution, TR of 4100ms, TE of 88ms, multi-band acceleration of 3 (Moeller et al., 2010), and a FOV of 240×240. The protocol used 6/8 partial Fourier acquisition in phase direction.

#### 2.2.2 Philips scanners

Matrix, FOV, and resolution parameters were identical across Philips and Siemens scanners in sMRI sessions. The T1-weighted session employed 225 slices, TR of 6.31ms, and TE of 2.9ms; the T2-weighted session used 256 slices, TR of 2500ms, and TE of 251.6ms. dMRI sessions shared similar slices, FOV, resolution, and multi-band acceleration factors, with variations in matrix (140 × 141). Philips protocol had longer TR (5300ms) and TE (89ms). The protocol used 0.6 partial Fourier acquisition in phase direction.

#### 2.2.3 GE scanners

GE sMRI sessions mirrored Siemens sessions in matrix, FOV, resolution, and TR, with 208 slices and TE of 2ms for T1-weighted and TE of 60ms for T2-weighted. GE dMRI sessions slightly differed from Siemens dMRI sessions in TE (81.9ms). The protocol used 5.5/8 partial Fourier acquisition in phase direction.

### 2.3 Imaging data preprocessing

#### 2.3.1 ABCD sMRI and dMRI preprocessing

The details of the preprocessing pipelines from ABCD Data Analysis and Informatics Core is described elsewhere(Hagler, Hatton et al. 2019). Briefly, for sMRI scans, this pipeline performs gradient distortion correction(Wald, Schmitt et al. 2001, Jovicich, Czanner et al. 2006) using scanner-specific and non-linear transformations provided by each scanner manufacturer, and intensity inhomogeneity correction(Sled, Zijdenbos et al. 1998, Ashburner and Friston 2000). T2-weighted images are then registered to T1-weighted images by maximizing mutual information(Wells, Viola et al. 1996), and resampled with a 1.0mm isotropic resolution reference brain in standard space. Subsequently, cortical reconstruction was performed using FreeSurfer 5.3.0, and reconstructed cortical surfaces were then registered to the Desikan atlas(Desikan, Ségonne et al. 2006). Average cortical thickness (CT) within each fuzzy-cluster parcels was calculated using smoothed surface maps (Chen, Gutierrez et al. 2012).

dMRI scans underwent eddy current correction using a diffusion gradient and model-based approach (Zhuang, Hrabe et al. 2006). Dark slices affected by abrupt head motion were identified through robust tensor fitting, and frames exceeding a normalized residual error threshold were censored. Head motion correction was performed by rigid-body-registering each frame to the corresponding volume synthesized from the post-ECC censored tensor fit. Gradient nonlinearity distortions were also corrected for each frame. The dMRI images were registered to T1-weighted structural images using mutual information (Wells, Viola et al. 1996), resampled with 1.7mm isotropic resolution, and adjusted for head rotation to achieve consistent diffusion orientations across participants.

#### 2.3.2 ABCD regional FA analysis

Regional FA values were extracted for major white matter fiber tracts using AtlasTrack approach that included fitting of the diffusion tensor model to the pre-processed diffusion images (Hagler, Ahmadi et al. 2008). Diffusion tensor imaging (DTI) measures, including fractional anisotropy (FA) were obtained using a standard linear estimation approach, with two different tensor model fits: one excluding frames with b>1000 s/mm^2^ (DTI inner shell) and another including all gradient strengths/shells (Alexander, Lee et al. 2007). In this study, we analyzed ABCD FA data extracted the full shell white matter FA modality.

#### 2.3.3 Final sample composition

The ABCD study release 5.0 includes the baseline scans from 11,868 subjects at 21 sites. The follow up scans included: N=3360 participants underwent one assessment only; 5744 participants underwent two assessments; 2619 participants completed all three assessments. After implementing the QC and preprocessing pipelines, the ABCD study provided brain white matter FA data for 11,542 subjects (mean baseline age 9.9 years [SD 0.6]; 47.9% female) and morphometric measures (i.e., CT) for 11,802 subjects (mean baseline age 9.9 years [SD 0.6]; 47.8% female). Subjects with only one assessment were also excluded, as longitudinal information is imperative for the evaluation of imaging data quality in our proposed method. 97 participants who went to different study sites during the baseline visit and follow-up visits were also excluded from the study. The distribution of subjects with multiple assessments across the 21 study sites is summarized in **Table S1** in the supplementary material. There are 7,889 subjects with FA data (mean baseline age 9.9 years [SD 0.6]; 46.6% female) and 8,326 with CT data (mean baseline age 9.9 years [SD 0.6]; 46.5% female).

### 2.4 Reliability analysis

In reliability analysis of neuroimaging data, the commonly used test-retest reliability scores are often built on the contrast of intra-subject vs inter-subject variances. A commonly used rationale is that the reliability is higher when the proportion of intra-subject variance is lower. The underlying assumption for this rationale is that each subject is measured repeatedly in the same condition (or with very high similarity). However, this assumption may not be applied to neuroimaging data that are subject to the rapid brain development of the adolescents, where intra-subject changes are assumed and the relatively lower or lack of intra-subject changes in specific tracts or individuals may even infer neurodevelopment abnormality. Therefore, the contribution of intra-subject variability to reliability estimate under typical ICC is inflated, and age correction is required for unbiased estimation of testing-retesting measures. Due to the similar enrollment age across sites, the population-level age-related brain (imaging measures) developmental trajectories is relatively invariant across the 21 sites, allowing us to integrate data from all sites to estimate the age trajectories. Specifically, we calculate the AICC as follows:

Step 1: Fit a weighted mixed model across all-sites to estimate the age trajectory parameters.

Step 2: For each site, perform age-correction using the estimated age trajectory parameters and calculate ICC. Update the weights based on the site-wise ICC values.

Step 3: Repeat step 1 and 2 until convergence and report the updated AICC values for all sites.

In step 1, the weighted mixed model is expressed as:

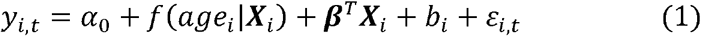

where *y*_*i*,*t*_ is the outcome (e.g., FA value on a white matter fiber tract, *i* denotes the subject (*i* = 1,…., *n*) and *t* denotes the time point; *f*(*age*_*i*_ |***X***_*i*_ ) is function of developmental trajectory by aging and trajectory can be sex and racial/ethnical group specific; ***X***_*i*_ is the vector of demographic variables; *b*_*i*_ is the random effect. The random slopes are generally not used because they tend to give rise to the convergence issue due to singular fit (i.e., overfit) (José C. Pinheiro 2024). The weight is incorporated in the covariance matrix 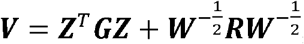, where ***Z*** is the design matrix giving the values of random effects to each observation and ***G*** is the covariance for the random effects and ***R*** denotes the dependence between repeated measures. The weight {*w*_*i*,*t*_} is calculated in step 2 as the input.

In step 2, we first perform ICC for each site. The weights are calculated using min-max scaling: 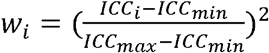, *ICC*_*i*_ is the ICC for the site where participant *i* was collected. By repeating steps 1 and 2, *f*(*age*_*i*_ |***X***_*i*_) can be estimated based on all participants while not being affected by the measurement errors. In practice, AICC converges fast. Since AICC is built on the mixed model, it is robust to drop-outs. Extensive simulation analysis was performed, and the results showed that AICC can estimate the underlying ICC more accurately than classical testing-testing measures. Details of the simulation study are included in **SI.1** in the supplementary material.

## 3. Results

### Reliability in CT and FA

The AICC was higher for the whole-brain gray matter CT measurements (AICC= 0.76±0.12), followed by whole-brain FA (0.61± 0.19). The AICC measurements per site are presented in **Figure 1a** and in **Table S2** from the supplementary material. Sites that used Siemens scanners showed higher AICC on average than GE and Philips for both FA (0.71±0.12 vs 0.46±0.17) and CT (0.80±0.10 vs 0.69±0.11). Both differences were significant (p < 0.001) (**Figure 1b**).

**Figure 1.**
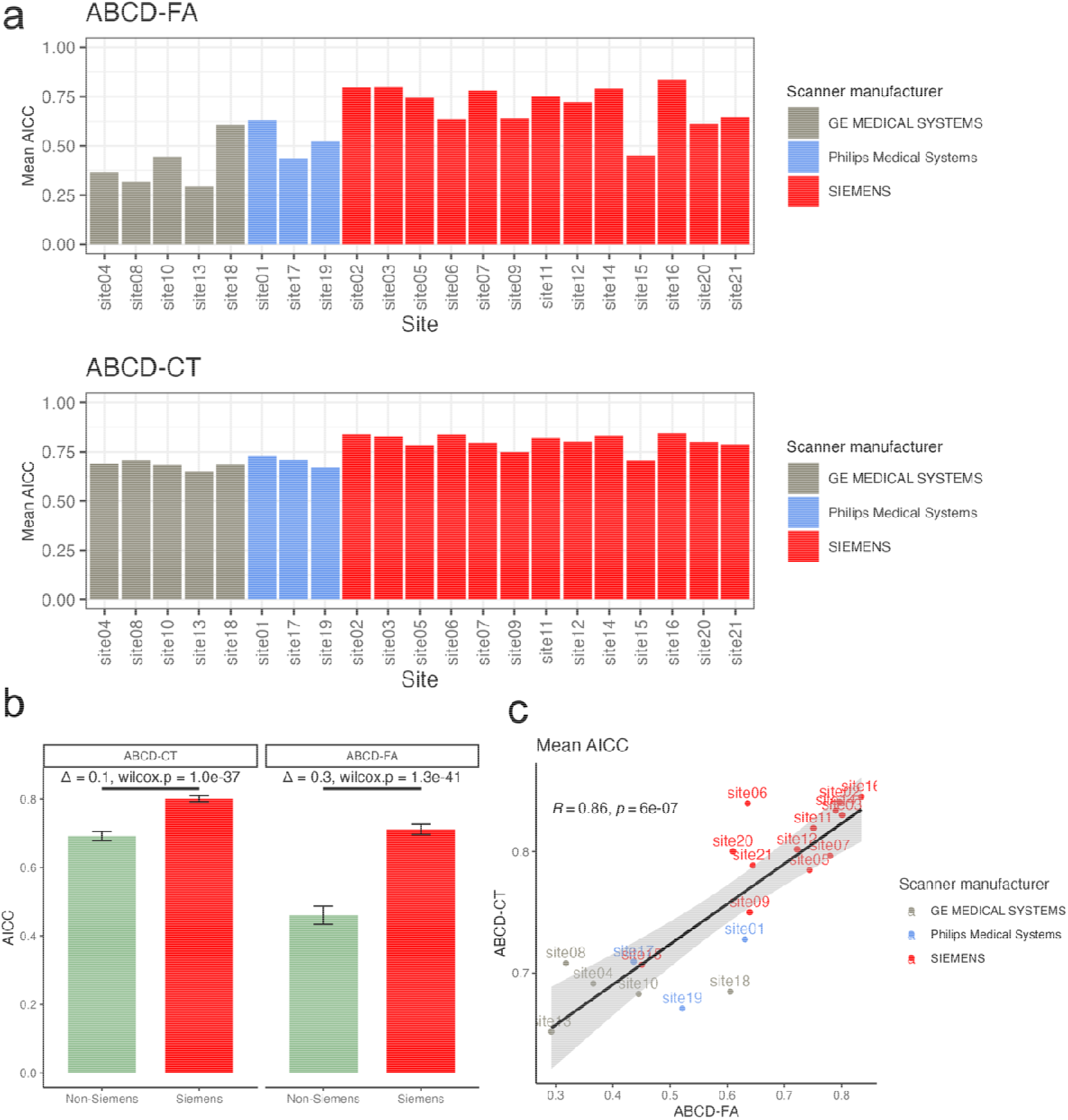
Mean AICC across brain regions for each site. The bar plots demonstrate the mean data reliability (i.e. AICC) of each site in the ABCD study. *Red* bars indicate sites using Siemens scanners, *blue* Philips, and *grey* GE. Panel (a) shows the mean AICC across all brain regions for FA and CT, respectively. Overall, data reliability was better in Siemens sites for both FA (AICC=0.71±0.12 vs 0.46±0.17, p<0.001) and CT (AICC=0.80±0.10 vs 0.69±0.11, p<0.001). Morphometry measures (i.e., CT) has higher reliability (AICC=0.76±0.12) than FA measures (AICC=0.61± 0.19) and less variations across sites-and-scanners. Panel (b) displays the significantly higher AICC of FA and CT from Siemens sites compared to non-Siemens sites. The error-bars indicate standard errors across sites and regions. Panel (c) shows the correlation of mean AICC between the two measures. Linear regression analysis showed a significant and positive correlation (r=0.86, p<0.001). This correlation was primarily driven by differences in scanners.

The site-wise AICC measurements for CT and FA showed significant and positive correlation, that is, sites with higher AICC for CT also had higher AICC for FA (r=0.86, p<0.001) (**Figure 1c**). This correlation was driven by differences in scanners – partial correlation adjusting for scanner manufacturers dropped to 0.62 with p=0.005.

### Regional differences

Regional differences in AICC for FA and CT are shown in **Figure 2** and **Table S3-4**. For FA values, the highest AICC values were observed for the Superior Cortico Striatal ∼0.75 (SD 0.16). The lowest AICC were observed in Forceps Minor and Fornix (excluding fimbria) with a mean value of 0.49 (SD 0.17) and 0.53 (SD 0.28), respectively. The regional AICC for CT were lowest for Cingulate and Para hippocampal gyri and highest for temporal pole and insula.

**Figure 2.**
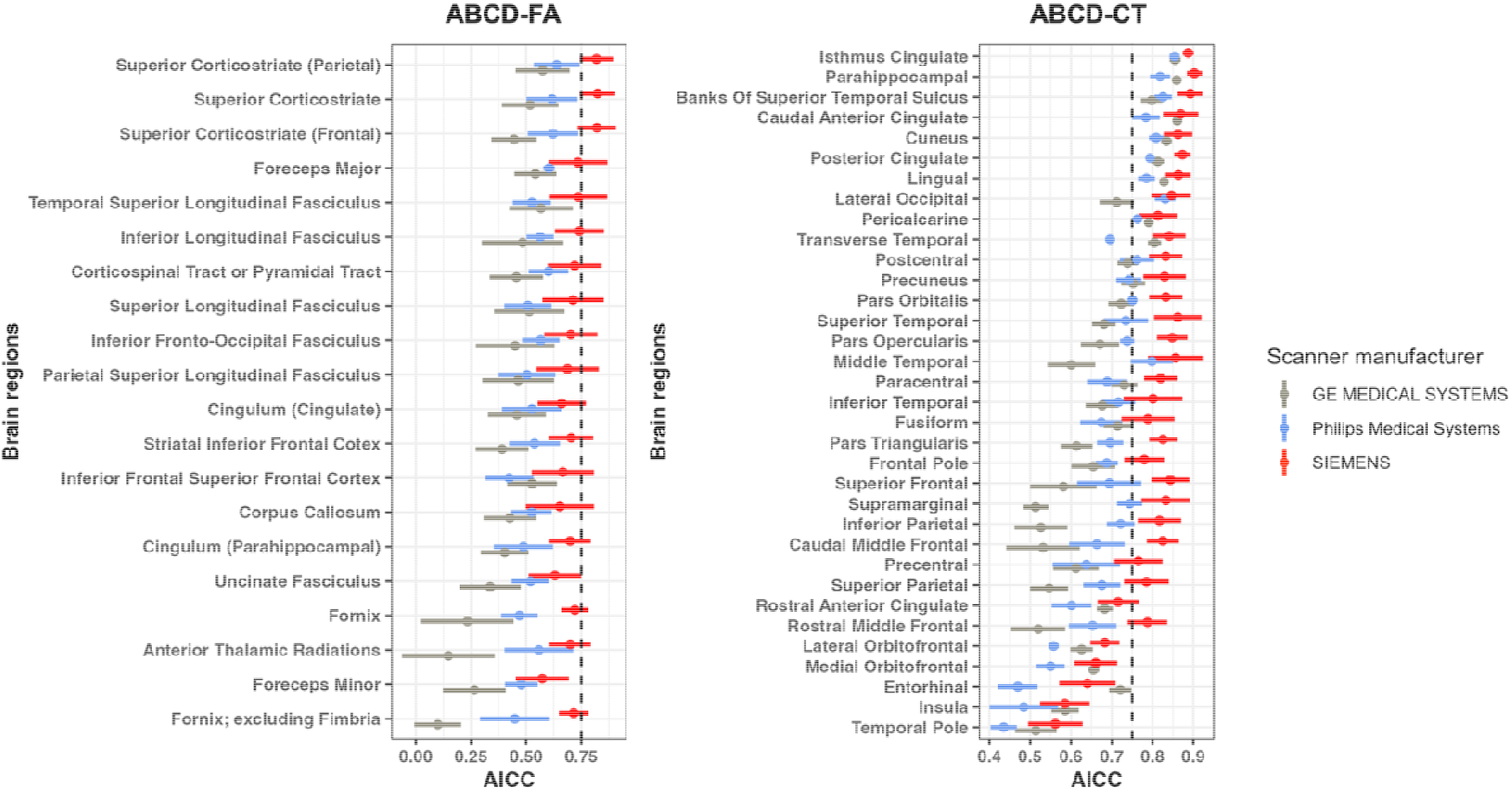
Regional differences of the mean AICC across sites, grouped by the type of scanners used. The forest plots represent the mean regional AICC across sites using scanners from the same manufacturer, with error bars indicating standard deviation. *Red* color shows the measures obtained from Siemens scanners, *blue* Philips, and *grey* GE. The dotted line is at the conventionally good ICC score, 0.75. Overall, for FA, Siemens sites have the highest AICC with small variations across all the individual tracts among the three types of scanners. For CT, differences in AICC related to scanners are notably smaller compared to both FA measures. Nevertheless, the temporal pole, insula, and entorhinal cortex consistently exhibited low mean AICC across sites for CT, regardless of the scanner type employed.

### Age trajectories

For Siemens scanners, the whole-brain FAs showed significant correlation with age (r=0.38, p<0.001) (**Figure 3**). Data collected from GE and Philips scanners showed lower correlations (FA: r=-0.05, p<0.001). CT demonstrated higher reliability across all sites and show smaller differences between the trajectories obtained from Siemens (r=-0.35, p<0.001) and non-Siemens sites (r=-0.31, p<0.001), but the effect size of the correlation was still attenuated in non-Siemens sites.

**Figure 3.**
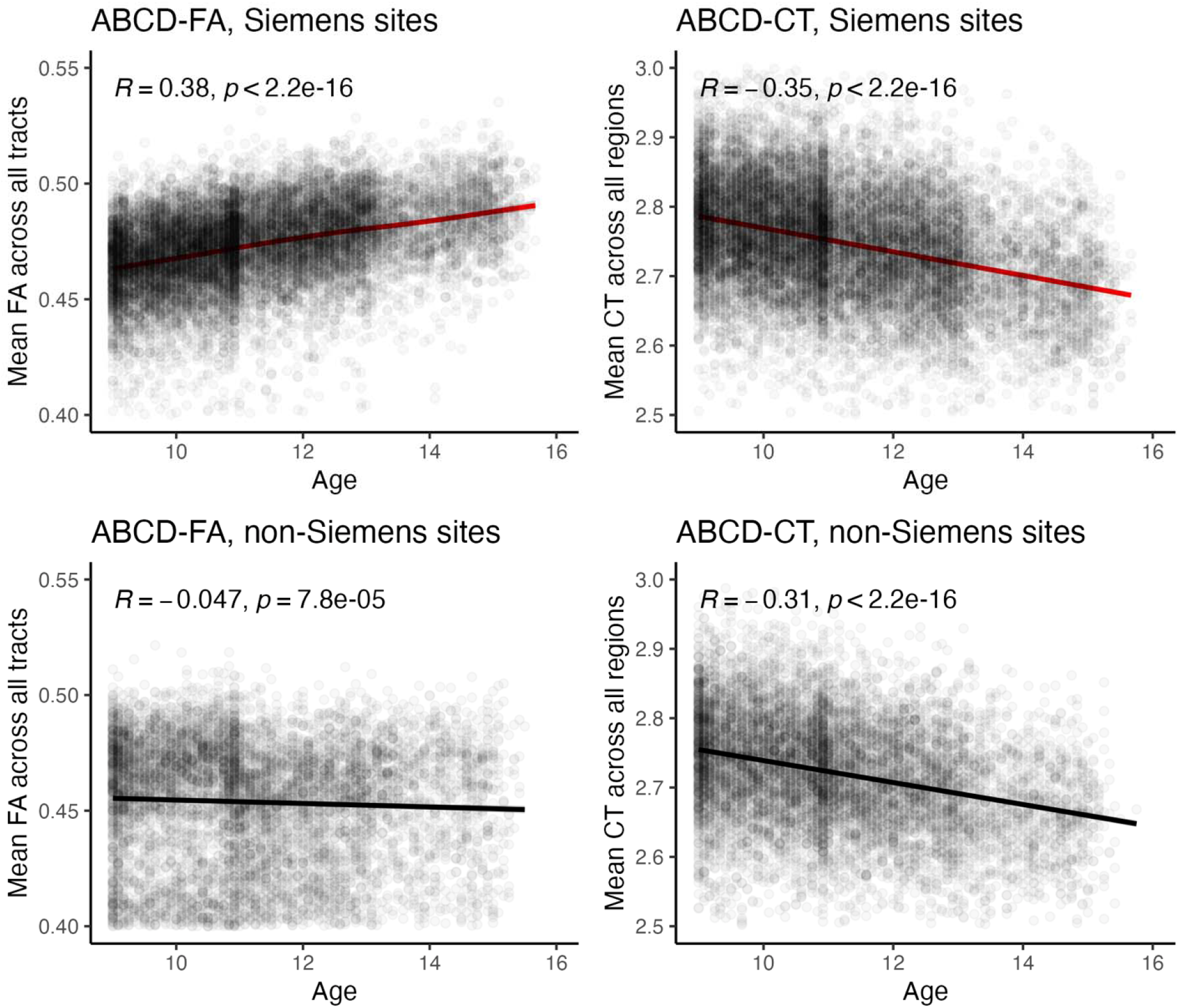
Correlation between age and imaging outcomes. The correlations between age and FA are positive in Siemens sites, indicating a myelination of WM in development as expected (r=0.38, p<0.001), while the age effect is reversed in non-Siemens sites (r=-0.05, p<0.001). The differences in the correlations between CT values obtained from different scanners are smaller (Siemens sites: r=-0.35, p<0.001 vs. non-Siemens sites: r=-0.31, p<0.001).

## 4. Discussion

ABCD is the largest longitudinal cohort tracking adolescent brain development using standardized protocols for multi-site data collection. We examined reproducibility of fractional anisotropy values extracted using the standard ABCD diffusion tensor imaging (DTI) workflows and that of cortical thickness derived from the ABCD structural MRI pipelines. We analyzed neuroimaging data collected over 5-year period and therefore expected that maturational change in FA and CT. We developed a novel iterative ICC approach that considers population-level developmental trends. We showed that despite the great efforts in ensuring consistent data collection across Siemens, Philips and GE scanners, the scanner-manufacturer-related variations were large, with data collected on Philips and GE platforms showing significantly poorer test-retest reliability. We also showed that this manufacturer-specific reproducibility difference was reflected in both diffusion and cortical thickness data. The scanner-related data differences in ABCD structural and functional data have been reported before (Nielson, Pereira et al. 2018, Sinha and Raamana 2023), although they focused on baseline scans only but here we provided a quantification of reliability using longitudinal data. We speculate on two likely causes for this scanner-related issues across these analyses.

The first likely source of non-biological variance that lowered the reproducibility metrics for GE and Philips scanners may include possible protocol deviation and differences in machine hardware and software technology that cannot be homogenized across the manufacturers. The deviations from prescribed protocol parameters are more pervasive for data collected on Philips and GE versus Siemens scanners (Sinha and Raamana 2023). The deviation can also be caused by operator errors. For example, the GE MR750 software may not display all slices for DTI sequence and to overcome this, operator must change the prescribed TR parameter and then change it back to the protocol mandated value. This can introduce a human error, compared to Siemens platform that does not require this step. The deviation of the protocol can also be caused by the differences in the automated scanner optimization algorithms built in by manufacturers that lead to changes in critical sequence parameters such as echo and repetition time, unbeknown to the operator (Sinha and Raamana 2023). The second source of the scanner related difference may arise from the overall stability of the magnet and gradient systems, differences in the k-space sampling trajectories and others. For example, Philips and GE scanners used by ABCD have slower gradient slew rates and therefore longer echo spacing. This led the ABCD protocol to use more aggressive partial Fourier imaging. Siemens DTI protocol collected 75% of the k-space versus 69% for GE and only 55% for Philips. The slower gradient performance forced the DTI protocol for Philips scanners to be split into two parts that are concatenated at the analysis stage. This was not the case for Siemens and GE scanners where all data were collected in a single step. The default direction in which k-space is transversed is also different between Siemens and GE and Philips scanners. These differences are hard to harmonize and likely cause complex interaction with the multi-slice excitation, leading to differences in signal, noise, distortions, and Nyquist ghost locations that reduce the overall reproducibility.

We observed significant scanner effects in the by-region AICC analysis with data collected using Siemens scanners showing uniformly higher AICC across all sites. The regional differences in AICC were also in agreement with reproducibility study of FA values derived from the Enhancing Neuro Imaging Genetic Meta Analyses (ENIGMA) DTI pipeline (Acheson, Wijtenburg et al. 2017). Specifically, lower reproducibility of FA values in fornix tracts were reported in two independent cohorts: adolescents and adults. The Fornix is a long and narrow bundle that is hard to correct for individual anatomical variability (Acheson, Wijtenburg et al. 2017). Additionally, another reproducibility study on multimodal MRI brain phenotypes in youth subjects reported consistent results: the ICC of FA in Forceps Minor was the lowest (0.76, 95%CI [0.61, 0.86]), followed by left Uncinate Fasciculus (ICC=0.78, 95%CI [0.65, 0.87]). This implies that the low reproducibility observed in these tracts is likely to be attributable to technical aspects of FA measurements including alignment-related methodological confounds (e.g., misalignments in registration or partial voluming effects) rather than a failure to account for the developmental trajectory (M, FB et al. 2014). Furthermore, CT measurements demonstrated consistently low AICC regions within the temporal lobe (i.e. temporal pole, insula, entorhinal cortex) and in the medial/lateral orbitofrontal lobe. Seiger et. al’s work showed that these regions have the largest difference in region-wise CT estimations using two different methods, suggesting these regions are susceptible to methodological confounds when estimating CT values (Seiger, Ganger et al. 2018). The orbitofrontal region and temporal poles are commonly vulnerable tracts that may suffer from low reproducibility if the signal-to-noise ratio is low (Farrell, Landman et al. 2007, Mac Donald, Johnson et al. 2011, Shahim, Holleran et al. 2017).

Scanner platform had a large impact on the measured effects of biological age on longitudinal FA values in ABCD. During adolescent development, longitudinal rise in FA is hypothesized to indicate ongoing myelination of cerebral white matter(Kochunov, Glahn et al. 2011, Kochunov, Glahn et al. 2011). This trend was readily observed for FA collected using Siemens scanners. However, data collected on GE or Philips scanners showed a weak and negative association between FA and age. MRI-based cortical thickness is also expected to show developmental change. The ongoing myelination of white matter adjacent to cortical ribbon and pruning of cortical neurons leads to reduction of measured CT during adolescence(Kochunov, Glahn et al. 2011, Kochunov, Glahn et al. 2011). The negative trend in CT values was again readily detected in the data collected across Siemens, and the age effect observed in data collected from non-Siemens sites was also attenuated. The smaller age effect difference of CT between Siemens and non-Siemens sites can be explained by lower across-site variation in AICC and reduced disparity in measurement errors between Siemens and non-Siemens sites. These results underscore the substantial influence of imaging data reliability on association analyses and the credibility of neurobiological findings. The AICC of FA values from GE or Philips scanners are significantly lower compared to FA from Siemens scanners and CT from all scanners, indicating greater measurement errors. These elevated measurement errors could obscure the association between FA values and age. Therefore, a thorough quality assessment should be conducted prior to the primary analysis to ensure the reliability and replicability of the results.

The presence of measurement errors in neuroimaging data introduces biases in association analyses. Measurement error in imaging predictors can attenuate or weaken observed brain-behavior effect sizes, while measurement error in imaging outcomes can inflate estimates of standard errors (Kenny 1979). In the context of longitudinal data analysis, non-negligible measurement errors can also cause the ordinary maximum likelihood estimators to be inconsistent (Fuller 2009). To address measurement errors, potential strategies involve careful consideration of data quality based on both data-driven methods such as AICC and manual quality control (MQC) measures. Quality assessment using the MQC method on the ABCD T1-weighted images revealed that 55.1% of the scans were identified as lower-quality images, contributing to bias in the statistical analysis outcomes (Elyounssi, Kunitoki et al. 2023). Therefore, the evaluation of data reliability becomes crucial for the robustness and accuracy of longitudinal multimodal imaging data analysis. While MQC is a valuable method for flexible quality assessment, its inherent labor-intensive nature poses practical challenges, particularly in the context of large-scale studies. In such scenarios, quantitative data consistency metrics like AICC can offer a more efficient and scalable alternative, allowing for data quality assessments with enhanced speed and consistency.

Ensuring high data quality is critical for large-scale studies seeking meaningful insights from imaging data analysis. Both correction methods and rigorous quality control filtering are essential for reducing measurement errors. Excluding subjects or even sites with lower data quality decreases the sample size, while maintaining an optimal sample size could result in latent bias. Therefore, the challenge lies in striking a balance between sample size and noise. Towards these goals, we have devised a novel longitudinal reliability statistical method. This approach takes into account the expected normal age-related intra-subject variance, a crucial consideration to maximize the value of neurodevelopmental data. We identified critical scanner-type-related confounds for imaging data, particularly longitudinal diffusion imaging data collected during rapid neurodevelopment, and have recommended at least partial solutions and attention to the use of data from different sites for different types of data use.

## Data Availability

Data used in this study are publicly accessible from the Adolescent Brain and Cognitive Development (ABCD) study (https://abcdstudy.org) in the NIMH Data Archive (NDA).

## Reference

Acheson, A., S. Wijtenburg, L. Rowland, A. Winkler, C. W. Mathias, L. Hong, N. Jahanshad, B. Patel, P. Thompson, S. McGuire, P. Sherman, P. Kochunov and D. D. Dougherty (2017). “Reproducibility of tract-based white matter microstructural measures using the ENIGMA-DTI protocol.” Genes Brain Behav 7(2): 10.1002/brb1003.1615.

Alexander, A. L., J. E. Lee, M. Lazar and A. S. Field (2007). “Diffusion tensor imaging of the brain.” Neurotherapeutics 4(3): 316–329.

Ashburner, J. and K. J. Friston (2000). “Voxel-based morphometry--the methods.” NeuroImage 11(6 Pt 1): 805-821.

Barnea-Goraly, N., V. Menon, M. Eckert, L. Tamm, R. Bammer, A. Karchemskiy, C. C. Dant and A. L. Reiss (2005). “White Matter Development During Childhood and Adolescence: A Cross-sectional Diffusion Tensor Imaging Study.” Cerebral Cortex 15(12): 1848–1854.

Barnhart, H. X., M. J. Haber and L. I. Lin (2007). “An Overview on Assessing Agreement with Continuous Measurements.” Journal of Biopharmaceutical Statistics 17(4): 529–569.

Bartzokis, G., P. H. Lu, K. Tingus, M. F. Mendez, A. Richard, D. G. Peters, B. Oluwadara, K. A. Barrall, J. P. Finn, P. Villablanca, P. M. Thompson and J. Mintz (2010). “Lifespan trajectory of myelin integrity and maximum motor speed.” Neurobiology of aging 31(9): 1554–1562.

Basser, P. J. (1994). “Focal magnetic stimulation of an axon.” IEEE Transactions on Biomedical Engineering 41(6): 601–606.

Beaulieu, C. (2002). “The basis of anisotropic water diffusion in the nervous system -a technical review.” NMR Biomed 15(7-8): 435–455.

Budde, M. D., J. H. Kim, H. F. Liang, R. E. Schmidt, J. H. Russell, A. H. Cross and S. K. Song (2007). “Toward accurate diagnosis of white matter pathology using diffusion tensor imaging.” Magn Reson Med 57(4): 688–695.

Casey, B. J., T. Cannonier, M. I. Conley, A. O. Cohen, D. M. Barch, M. M. Heitzeg, M. E. Soules, T. Teslovich, D. V. Dellarco, H. Garavan, C. A. Orr, T. D. Wager, M. T. Banich, N. K. Speer, M. T. Sutherland, M. C. Riedel, A. S. Dick, J. M. Bjork, K. M. Thomas, … and A. M. Dale (2018). “The Adolescent Brain Cognitive Development (ABCD) study: Imaging acquisition across 21 sites.” Developmental Cognitive Neuroscience 32: 43–54.

Casey, B. J., R. M. Jones and T. A. Hare (2008). “The Adolescent Brain.” Annals of the New York Academy of Sciences 1124: 111–126.

Casey, B. J., J. T. Nigg and S. Durston (2007). “New potential leads in the biology and treatment of attention deficit-hyperactivity disorder.” Curr Opin Neurol 20(2): 119–124.

Chen, C.-H., E. D. Gutierrez, W. Thompson, M. S. Panizzon, T. L. Jernigan, L. T. Eyler, C. Fennema-Notestine, A. J. Jak, M. C. Neale, C. E. Franz, M. J. Lyons, M. D. Grant, B. Fischl, L. J. Seidman, M. T. Tsuang, W. S. Kremen and A. M. Dale (2012). “Hierarchical Genetic Organization of Human Cortical Surface Area.” Science (New York, N.Y.) 335(6076): 1634–1636.

Conturo, T. E., R. C. McKinstry, E. Akbudak and B. H. Robinson (1996). “Encoding of anisotropic diffusion with tetrahedral gradients: a general mathematical diffusion formalism and experimental results.” Magn Reson Med 35(3): 399–412.

Desikan, R. S., F. Ségonne, B. Fischl, B. T. Quinn, B. C. Dickerson, D. Blacker, R. L. Buckner, A. M. Dale, R. P. Maguire, B. T. Hyman, M. S. Albert and R. J. Killiany (2006). “An automated labeling system for subdividing the human cerebral cortex on MRI scans into gyral based regions of interest.” NeuroImage 31(3): 968–980.

Elyounssi, S., K. Kunitoki, J. A. Clauss, E. Laurent, K. Kane, D. E. Hughes, C. E. Hopkinson, O. Bazer, R. F. Sussman, A. E. Doyle, H. Lee, B. Tervo-Clemmens, H. Eryilmaz, R. L. Gollub, D. M. Barch, T. D. Satterthwaite, K. F. Dowling and J. L. Roffman (2023). “Uncovering and mitigating bias in large, automated MRI analyses of brain development.” bioRxiv: 2023.2002.2028.530498.

Farrell, J. A. D., B. A. Landman, C. K. Jones, S. A. Smith, J. L. Prince, P. C. M. v. Zijl and S. Mori (2007). “Effects of signal □ to □ noise ratio on the accuracy and reproducibility of diffusion tensor imaging–derived fractional anisotropy, mean diffusivity, and principal eigenvector measurements at 1.5T.” Journal of Magnetic Resonance Imaging 26(3).

Flechsig, P. (1901). “Developmental (myelogenetic) localisation of the cerebral cortex in the human.” Lancet 158: 1027–1030.

Fuller, W. A. (2009). Measurement Error Models, John Wiley & Sons.

Garavan, H., H. Bartsch, K. Conway, A. Decastro, R. Z. Goldstein, S. Heeringa, T. Jernigan, A. Potter, W. Thompson and D. Zahs (2018). “Recruiting the ABCD sample: Design considerations and procedures.” Developmental Cognitive Neuroscience 32: 16–22.

Gogtay, N., J. N. Giedd, L. Lusk, K. M. Hayashi, D. Greenstein, A. C. Vaituzis, T. F. Nugent, 3rd, D. H. Herman, L. S. Clasen, A. W. Toga, J. L. Rapoport and P. M. Thompson (2004). “Dynamic mapping of human cortical development during childhood through early adulthood.” Proc Natl Acad Sci U S A 101(21): 8174–8179.

Gogtay, N. and P. M. Thompson (2009). “Mapping gray matter development: implications for typical development and vulnerability to psychopathology.” Brain Cogn 72(1): 6–15.

Hagler, D. J., M. E. Ahmadi, J. Kuperman, D. Holland, C. R. McDonald, E. Halgren and A. M. Dale (2008). “Automated white□matter tractography using a probabilistic diffusion tensor atlas: Application to temporal lobe epilepsy.” Human Brain Mapping 30(5): 1535–1547.

Hagler, D. J., S. N. Hatton, M. D. Cornejo, C. Makowski, D. A. Fair, A. S. Dick, M. T. Sutherland, B. J. Casey, D. M. Barch, M. P. Harms, R. Watts, J. M. Bjork, H. P. Garavan, L. Hilmer, C. J. Pung, C. S. Sicat, J. Kuperman, H. Bartsch, F. Xue, … and A. M. Dale (2019). “Image processing and analysis methods for the Adolescent Brain Cognitive Development Study.” NeuroImage 202: 116091.

José C. Pinheiro , D. M. B. (2024). Mixed-Effects Models in S and S-PLUS.

Jovicich, J., S. Czanner, D. Greve, E. Haley, A. van der Kouwe, R. Gollub, D. Kennedy, F. Schmitt, G. Brown, J. Macfall, B. Fischl and A. Dale (2006). “Reliability in multi-site structural MRI studies: effects of gradient non-linearity correction on phantom and human data.” NeuroImage 30(2): 436–443.

Kalia, M. (2008). “Brain development: anatomy, connectivity, adaptive plasticity, and toxicity.” Metabolism 57 Suppl 2: S2-5.

Karcher, N. R. and D. M. Barch (2021). “The ABCD study: understanding the development of risk for mental and physical health outcomes.” Neuropsychopharmacology 46(1): 131–142.

Kenny, D. A. (1979). Correlation and causality.

Kochunov, P., D. Glahn, T. Nichols, A. Winkler, E. Hong, H. Holcomb, J. Stein, P. Thompson, J. Curran, M. Carless, R. Olvera, M. Johnson, S. Cole, V. Kochunov, J. Kent and J. Blangero (2011). “Genetic Analysis of Cortical Thickness and Fractional Anisotropy of Water Diffusion in the Brain.” Frontiers in Neuroscience 5(120): 1–15.

Kochunov, P., D. C. Glahn, J. Lancaster, P. M. Thompson, V. Kochunov, B. Rogers, P. Fox, J. Blangero and D. E. Williamson (2011). “Fractional anisotropy of cerebral white matter and thickness of cortical gray matter across the lifespan.” Neuroimage 58(1): 41–49.

Kochunov, P., P. M. Thompson, A. Winkler, M. Morrissey, M. Fu, T. R. Coyle, X. Du, F. Muellerklein, A. Savransky, C. Gaudiot, H. Sampath, G. Eskandar, N. Jahanshad, B. Patel, L. Rowland, T. E. Nichols, J. R. O’Connell, A. R. Shuldiner, B. D. Mitchell and L. E. Hong (2015). “The common genetic influence over processing speed and white matter microstructure: Evidence from the Old Order Amish and Human Connectome Projects.” Neuroimage 125: 189–197.

Kochunov, P., D. E. Williamson, J. Lancaster, P. Fox, J. Cornell, J. Blangero and D. C. Glahn (2012). “Fractional anisotropy of water diffusion in cerebral white matter across the lifespan.” Neurobiol Aging 33(1): 9–20.

Konrad, K., C. Firk and P. J. Uhlhaas (2013). “Brain Development During Adolescence.” Deutsches Ärzteblatt International 110(25): 425–431.

M, B., L. FB, L. A, T. CM, B. GJ, S. B and M.-H. KH (2014). “Methodological considerations on tract-based spatial statistics (TBSS).” NeuroImage 100.

Mac Donald, C. L., A. M. Johnson, D. Cooper, E. C. Nelson, N. J. Werner, J. S. Shimony, A. Z. Snyder, M. E. Raichle, J. R. Witherow, R. Fang, S. F. Flaherty and D. L. Brody (2011). “Detection of Blast-Related Traumatic Brain Injury in U.S. Military Personnel.” New England Journal of Medicine 364(22).

Madler, B., S. A. Drabycz, S. H. Kolind, K. P. Whittall and A. L. MacKay (2008). “Is diffusion anisotropy an accurate monitor of myelination? Correlation of multicomponent T2 relaxation and diffusion tensor anisotropy in human brain.” Magn Reson Imaging 26(7): 874–888.

Nielson, D. M., F. Pereira, C. Y. Zheng, N. Migineishvili, J. A. Lee, A. G. Thomas and P. A. Bandettini (2018). “Detecting and harmonizing scanner differences in the ABCD study - annual release 1.0.” bioRxiv: 309260.

Pierpaoli, C. and P. J. Basser (1996). “Toward a quantitative assessment of diffusion anisotropy.” Magn Reson Med 36(6): 893–906.

Rapoport, J. L., A. Addington and S. Frangou (2005). “The neurodevelopmental model of schizophrenia: what can very early onset cases tell us?” Curr Psychiatry Rep 7(2): 81–82.

Ryan, M. C., P. Sherman, L. M. Rowland, S. A. Wijtenburg, A. Acheson, E. Fieremans, J. Veraart, D. S. Novikov, L. E. Hong, J. Sladky, P. D. Peralta, P. Kochunov and S. A. McGuire (2017). “Miniature pig model of human adolescent brain white matter development.” Journal of Neuroscience Methods 296: 99–108.

Ryan, M. C., P. Sherman, L. M. Rowland, S. A. Wijtenburg, A. Acheson, E. Fieremans, J. Veraart, D. S. Novikov, L. E. Hong, J. Sladky, P. D. Peralta, P. Kochunov and S. A. McGuire (2018). “Miniature pig magnetic resonance spectroscopy model of normal adolescent brain development “ Journal of Neuroscience Methods 296: 99–108.

Seiger, R., S. Ganger, G. S. Kranz, A. Hahn and R. Lanzenberger (2018). “Cortical Thickness Estimations of FreeSurfer and the CAT12 Toolbox in Patients with Alzheimer’s Disease and Healthy Controls.” Journal of Neuroimaging 28(5): 515–523.

Shahim, P., L. Holleran, J. H. Kim, D. L. Brody, P. Shahim, L. Holleran, J. H. Kim and D. L. Brody (2017). “Test-retest reliability of high spatial resolution diffusion tensor and diffusion kurtosis imaging.” Scientific Reports 2017 7:1 7(1).

Shrout, P. E. and J. L. Fleiss (1979). “Intraclass correlations: uses in assessing rater reliability.” Psychological Bulletin 86(2): 420–428.

Sinha, H. and P. R. Raamana (2023). “Solving the Pervasive Problem of Protocol Non-Compliance in MRI using an Open-Source tool <em>mrQA</em>.” bioRxiv: 2023.2007.2017.548591.

Sled, J. G., A. P. Zijdenbos and A. C. Evans (1998). “A nonparametric method for automatic correction of intensity nonuniformity in MRI data.” IEEE transactions on medical imaging 17(1): 87–97.

Song, S. K., S. W. Sun, W. K. Ju, S. J. Lin, A. H. Cross and A. H. Neufeld (2003). “Diffusion tensor imaging detects and differentiates axon and myelin degeneration in mouse optic nerve after retinal ischemia.” Neuroimage 20(3): 1714–1722.

Song, S. K., J. Yoshino, T. Q. Le, S. J. Lin, S. W. Sun, A. H. Cross and R. C. Armstrong (2005). “Demyelination increases radial diffusivity in corpus callosum of mouse brain.” Neuroimage 26(1): 132–140.

Ulug, A. M., P. B. Barker and P. C. van Zijl (1995). “Correction of motional artifacts in diffusion-weighted images using a reference phase map.” Magn Reson Med 34(3): 476–480.

Wald, L., F. Schmitt and A. Dale (2001). “Systematic spatial distortion in MRI due to gradient non-linearities.” NeuroImage 6 Supplement(13): 50.

Wells, W. M., P. Viola, H. Atsumi, S. Nakajima and R. Kikinis (1996). “Multi-modal volume registration by maximization of mutual information.” Medical Image Analysis 1(1): 35–51.

Wijtenburg, S. A., F. E. Gaston, E. A. Spieker, S. A. Korenic, P. Kochunov, L. E. Hong and L. M. Rowland (2013). “Reproducibility of phase rotation STEAM at 3T: focus on glutathione.” Magn Reson Med 72(3): 603–609.

Xue, C., J. Yuan, G. G. Lo, A. T. Y. Chang, D. M. C. Poon, O. L. Wong, Y. Zhou and W. C. W. Chu (2021). “Radiomics feature reliability assessed by intraclass correlation coefficient: a systematic review.” Quantitative Imaging in Medicine and Surgery 11(10): 4431–4460.

Zhuang, J., J. Hrabe, A. Kangarlu, D. Xu, R. Bansal, C. A. Branch and B. S. Peterson (2006). “Correction of Eddy-Current Distortions in Diffusion Tensor Images Using the Known Directions and Strengths of Diffusion Gradients.” Journal of magnetic resonance imaging : JMRI 24(5): 1188.

Zuo, X.-N. and X.-X. Xing (2014). “Test-retest reliabilities of resting-state FMRI measurements in human brain functional connectomics: A systems neuroscience perspective.” Neuroscience & Biobehavioral Reviews 45: 100–118.

