## supplementary material for "A site-wise reliability analysis of the ABCD diffusion fractional anisotropy and cortical thickness: impact of scanner platforms"

### Table S1. Distribution of subjects with multiple assessments in each site, stratified by imaging outcomes

| SITE ID | No. of subjects in each site | |
| --- | --- | --- |
|  | ABCD-FA | ABCD-CT |
| site01 | 226 | 237 |
| site02 | 426 | 442 |
| site03 | 356 | 374 |
| site04 | 609 | 646 |
| site05 | 272 | 277 |
| site06 | 395 | 408 |
| site07 | 212 | 220 |
| site08 | 153 | 187 |
| site09 | 223 | 232 |
| site10 | 538 | 593 |
| site11 | 263 | 276 |
| site12 | 394 | 409 |
| site13 | 450 | 481 |
| site14 | 450 | 462 |
| site15 | 237 | 256 |
| site16 | 743 | 764 |
| site17 | 366 | 384 |
| site18 | 271 | 292 |
| site19 | 326 | 358 |
| site20 | 582 | 614 |
| site21 | 397 | 414 |
| Total | 7889 | 8326 |

Table S2. Mean AICC (SD) across regions for each site, by imaging measures

| **Site** | **Scanner** | **ABCD-FA** | **ABCD-CT** |
| --- | --- | --- | --- |
| site01 | Philips Medical Systems | 0.63 (0.06) | 0.73 (0.1) |
| site02 | SIEMENS | 0.8 (0.05) | 0.84 (0.07) |
| site03 | SIEMENS | 0.8 (0.07) | 0.83 (0.08) |
| site04 | GE MEDICAL SYSTEMS | 0.39 (0.18) | 0.69 (0.12) |
| site05 | SIEMENS | 0.75 (0.08) | 0.78 (0.11) |
| site06 | SIEMENS | 0.64 (0.07) | 0.84 (0.08) |
| site07 | SIEMENS | 0.78 (0.07) | 0.79 (0.08) |
| site08 | GE MEDICAL SYSTEMS | 0.33 (0.11) | 0.71 (0.1) |
| site09 | SIEMENS | 0.64 (0.1) | 0.75 (0.1) |
| site10 | GE MEDICAL SYSTEMS | 0.46 (0.18) | 0.68 (0.11) |
| site11 | SIEMENS | 0.75 (0.06) | 0.82 (0.09) |
| site12 | SIEMENS | 0.72 (0.07) | 0.8 (0.08) |
| site13 | GE MEDICAL SYSTEMS | 0.31 (0.16) | 0.66 (0.13) |
| site14 | SIEMENS | 0.79 (0.07) | 0.83 (0.08) |
| site15 | SIEMENS | 0.46 (0.09) | 0.71 (0.09) |
| site16 | SIEMENS | 0.83 (0.05) | 0.84 (0.09) |
| site17 | Philips Medical Systems | 0.45 (0.07) | 0.71 (0.12) |
| site18 | GE MEDICAL SYSTEMS | 0.59 (0.13) | 0.69 (0.11) |
| site19 | Philips Medical Systems | 0.52 (0.09) | 0.67 (0.11) |
| site20 | SIEMENS | 0.61 (0.11) | 0.8 (0.1) |
| site21 | SIEMENS | 0.65 (0.07) | 0.79 (0.09) |

Table S3. Mean regional ABCD-FA AICC (SD) across sites, by scanner platforms

| **Tract** | **GE MEDICAL SYSTEMS** | **Philips Medical Systems** | **SIEMENS** |
| --- | --- | --- | --- |
| Banks Of Superior Temporal Sulcus | 0.8 (0.03) | 0.83 (0.02) | 0.89 (0.03) |
| Caudal Anterior Cingulate | 0.86 (0.01) | 0.78 (0.03) | 0.87 (0.04) |
| Caudal Middle Frontal | 0.53 (0.09) | 0.66 (0.07) | 0.83 (0.04) |
| Cuneus | 0.83 (0.01) | 0.81 (0.02) | 0.86 (0.04) |
| Entorhinal | 0.72 (0.03) | 0.47 (0.05) | 0.64 (0.07) |
| Frontal Pole | 0.65 (0.05) | 0.69 (0.03) | 0.78 (0.05) |
| Fusiform | 0.71 (0.03) | 0.67 (0.05) | 0.79 (0.07) |
| Inferior Parietal | 0.53 (0.06) | 0.72 (0.03) | 0.82 (0.05) |
| Inferior Temporal | 0.68 (0.04) | 0.72 (0.04) | 0.8 (0.07) |
| Insula | 0.59 (0.03) | 0.48 (0.08) | 0.59 (0.06) |
| Isthmus Cingulate | 0.86 (0.01) | 0.85 (0.01) | 0.89 (0.01) |
| Lateral Occipital | 0.71 (0.04) | 0.83 (0.03) | 0.85 (0.05) |
| Lateral Orbitofrontal | 0.63 (0.03) | 0.56 (0.01) | 0.68 (0.04) |
| Lingual | 0.83 (0.01) | 0.78 (0.02) | 0.86 (0.03) |
| Medial Orbitofrontal | 0.66 (0.01) | 0.55 (0.03) | 0.66 (0.05) |
| Middle Temporal | 0.6 (0.06) | 0.8 (0.05) | 0.86 (0.07) |
| Paracentral | 0.73 (0.03) | 0.69 (0.05) | 0.82 (0.04) |
| Parahippocampal | 0.86 (0) | 0.82 (0.02) | 0.9 (0.02) |
| Pars Opercularis | 0.67 (0.05) | 0.74 (0.02) | 0.85 (0.04) |
| Pars Orbitalis | 0.72 (0.03) | 0.75 (0.01) | 0.83 (0.04) |
| Pars Triangularis | 0.61 (0.04) | 0.7 (0.03) | 0.83 (0.03) |
| Pericalcarine | 0.79 (0.01) | 0.76 (0.01) | 0.81 (0.05) |
| Postcentral | 0.74 (0.02) | 0.76 (0.04) | 0.83 (0.04) |
| Posterior Cingulate | 0.81 (0.02) | 0.79 (0.01) | 0.87 (0.02) |
| Precentral | 0.61 (0.06) | 0.64 (0.08) | 0.77 (0.06) |
| Precuneus | 0.75 (0.03) | 0.74 (0.03) | 0.83 (0.05) |
| Rostral Anterior Cingulate | 0.68 (0.02) | 0.6 (0.05) | 0.72 (0.05) |
| Rostral Middle Frontal | 0.52 (0.07) | 0.65 (0.06) | 0.79 (0.05) |
| Superior Frontal | 0.58 (0.08) | 0.69 (0.08) | 0.84 (0.05) |
| Superior Parietal | 0.55 (0.05) | 0.68 (0.05) | 0.78 (0.06) |
| Superior Temporal | 0.68 (0.03) | 0.73 (0.05) | 0.86 (0.06) |
| Supramarginal | 0.51 (0.03) | 0.74 (0.03) | 0.83 (0.06) |
| Temporal Pole | 0.51 (0.05) | 0.44 (0.03) | 0.56 (0.07) |
| Transverse Temporal | 0.8 (0.02) | 0.7 (0.01) | 0.84 (0.04) |

Table S4. Mean regional ABCD-CT AICC (SD) across sites, by scanner platforms

| **Region** | **GE MEDICAL SYSTEMS** | **Philips Medical Systems** | **SIEMENS** |
| --- | --- | --- | --- |
| Banks Of Superior Temporal Sulcus | 0.8 (0.03) | 0.83 (0.02) | 0.89 (0.03) |
| Caudal Anterior Cingulate | 0.86 (0.01) | 0.78 (0.03) | 0.87 (0.04) |
| Caudal Middle Frontal | 0.53 (0.09) | 0.66 (0.07) | 0.83 (0.04) |
| Cuneus | 0.83 (0.01) | 0.81 (0.02) | 0.86 (0.04) |
| Entorhinal | 0.72 (0.03) | 0.47 (0.05) | 0.64 (0.07) |
| Frontal Pole | 0.65 (0.05) | 0.69 (0.03) | 0.78 (0.05) |
| Fusiform | 0.71 (0.03) | 0.67 (0.05) | 0.79 (0.07) |
| Inferior Parietal | 0.53 (0.06) | 0.72 (0.03) | 0.82 (0.05) |
| Inferior Temporal | 0.68 (0.04) | 0.72 (0.04) | 0.8 (0.07) |
| Insula | 0.59 (0.03) | 0.48 (0.08) | 0.59 (0.06) |
| Isthmus Cingulate | 0.86 (0.01) | 0.85 (0.01) | 0.89 (0.01) |
| Lateral Occipital | 0.71 (0.04) | 0.83 (0.03) | 0.85 (0.05) |
| Lateral Orbitofrontal | 0.63 (0.03) | 0.56 (0.01) | 0.68 (0.04) |
| Lingual | 0.83 (0.01) | 0.78 (0.02) | 0.86 (0.03) |
| Medial Orbitofrontal | 0.66 (0.01) | 0.55 (0.03) | 0.66 (0.05) |
| Middle Temporal | 0.6 (0.06) | 0.8 (0.05) | 0.86 (0.07) |
| Paracentral | 0.73 (0.03) | 0.69 (0.05) | 0.82 (0.04) |
| Parahippocampal | 0.86 (0) | 0.82 (0.02) | 0.9 (0.02) |
| Pars Opercularis | 0.67 (0.05) | 0.74 (0.02) | 0.85 (0.04) |
| Pars Orbitalis | 0.72 (0.03) | 0.75 (0.01) | 0.83 (0.04) |
| Pars Triangularis | 0.61 (0.04) | 0.7 (0.03) | 0.83 (0.03) |
| Pericalcarine | 0.79 (0.01) | 0.76 (0.01) | 0.81 (0.05) |
| Postcentral | 0.74 (0.02) | 0.76 (0.04) | 0.83 (0.04) |
| Posterior Cingulate | 0.81 (0.02) | 0.79 (0.01) | 0.87 (0.02) |
| Precentral | 0.61 (0.06) | 0.64 (0.08) | 0.77 (0.06) |
| Precuneus | 0.75 (0.03) | 0.74 (0.03) | 0.83 (0.05) |
| Rostral Anterior Cingulate | 0.68 (0.02) | 0.6 (0.05) | 0.72 (0.05) |
| Rostral Middle Frontal | 0.52 (0.07) | 0.65 (0.06) | 0.79 (0.05) |
| Superior Frontal | 0.58 (0.08) | 0.69 (0.08) | 0.84 (0.05) |
| Superior Parietal | 0.55 (0.05) | 0.68 (0.05) | 0.78 (0.06) |
| Superior Temporal | 0.68 (0.03) | 0.73 (0.05) | 0.86 (0.06) |
| Supramarginal | 0.51 (0.03) | 0.74 (0.03) | 0.83 (0.06) |
| Temporal Pole | 0.51 (0.05) | 0.44 (0.03) | 0.56 (0.07) |
| Transverse Temporal | 0.8 (0.02) | 0.7 (0.01) | 0.84 (0.04) |

SI.1 Simulation study

We performed simulation studies to evaluate the performance of our proposed AICC method and benchmarked it against age-corrected ICC using classic mixed effect models. First, we simulated imaging-derived phenotype data in a multi-site setting with 20 sites, resembling the ABCD study. We considered that a proportion 𝑝 of the sites contributed low-quality imaging data, while the remaining sites (1- 𝑝) reported more reliable data. In this setup, we assumed that reliable data followed a replicable developmental trajectory characterized by a linear age effect parameter $\beta$, while the low-quality data showed an attenuated or nullified age-affect $\beta^{*}$, as observed in the ABCD study^1^. The outcome $Y_{ij}\mathcal{\sim N}\left( \beta\times Age_{ij},\Sigma\left( \rho\right) \right)$ was generated for $i=1,\ldots n$ subjects and $j=1,2,3$ timepoints at $20\left( 1-p \right)$reliable sites independently, where $Age_{ij}$ represents the age of subject $i$ at timepoint $j$, and $\Sigma\left( \rho\right)=\sigma^{2}\left[ \begin{matrix} 1 & \rho& \rho\\ \rho& 1 & \rho\\ \rho& \rho& 1 \end{matrix} \right]$ is the covariance matrix with correlation ρ between each time point. By definition, $\rho$ is also equivalent to the age-corrected ICC of the outcome $Y_{ij}$ across all timepoints ^2,3^. Similarly, we generated the low-quality outcome $Y_{ij}^{*}\mathcal{\sim N}\left( \beta^{*}\times Age_{ij},\Sigma^{*}\left( \rho^{*} \right) \right)$ for $20p$ sites. We assumed that the low-quality imaging data can lead to 1) reduced reliability reflected by ICC and 2) biased age effect estimation.

$Age_{i}$ at baseline was generated using a uniform distribution ranging from 9 to 11 for all sites, and an age increment of 2 at each later time point. A sample size of $n=50$ were used for each site. We varied $p$ and $\beta$, i.e., $p\in\{0.1, 0.15, 0.2, \ldots, 0.85, 0.9\}$ and $\beta\in\{0.1, 0.3, 0.5\},$to simulate diverse overall data quality and effect sizes of brain development. We set $\rho=0.8$, $\rho^{*}= 0.4$, $\sigma^{2}=0.1$, and $\beta^{*}=0.5\beta$. Each parameter combination underwent 100 simulations. In each simulation, we performed the proposed AICC-estimation procedure on the generated data set. The estimation bias $\hat{\beta}-\beta$ was used as the evaluation criteria to compare the estimated age effects. Accuracy of the estimated testing-retesting reliability adjusting for the aging trend was also evaluated.

The results were summarized in Figure S1. Our method demonstrated consistent accuracy and robustness in estimating both age effects and testing-retesting reliability. The accuracy of the estimation was influenced by $p$, the proportion of contaminated data, and $\beta$, the true population age effect. The percent bias of the estimates increases as $p$ increases in the data set of interest; however, our method consistently outperforms the testing-retesting measure obtained using classic mixed models when $p$ was below 70%. Additionally, the AICC estimated by our method approached being unbiased when $p<50\%$, irrespective of the $\beta$ assumption. A notable advantage of our method lies in its reduced sensitivity to measurement errors within the dataset compared to conventional approaches. Consequently, it can accurately capture the testing-retesting reliability of longitudinally measured imaging data while accounting for age effects amidst the presence of measurement errors.

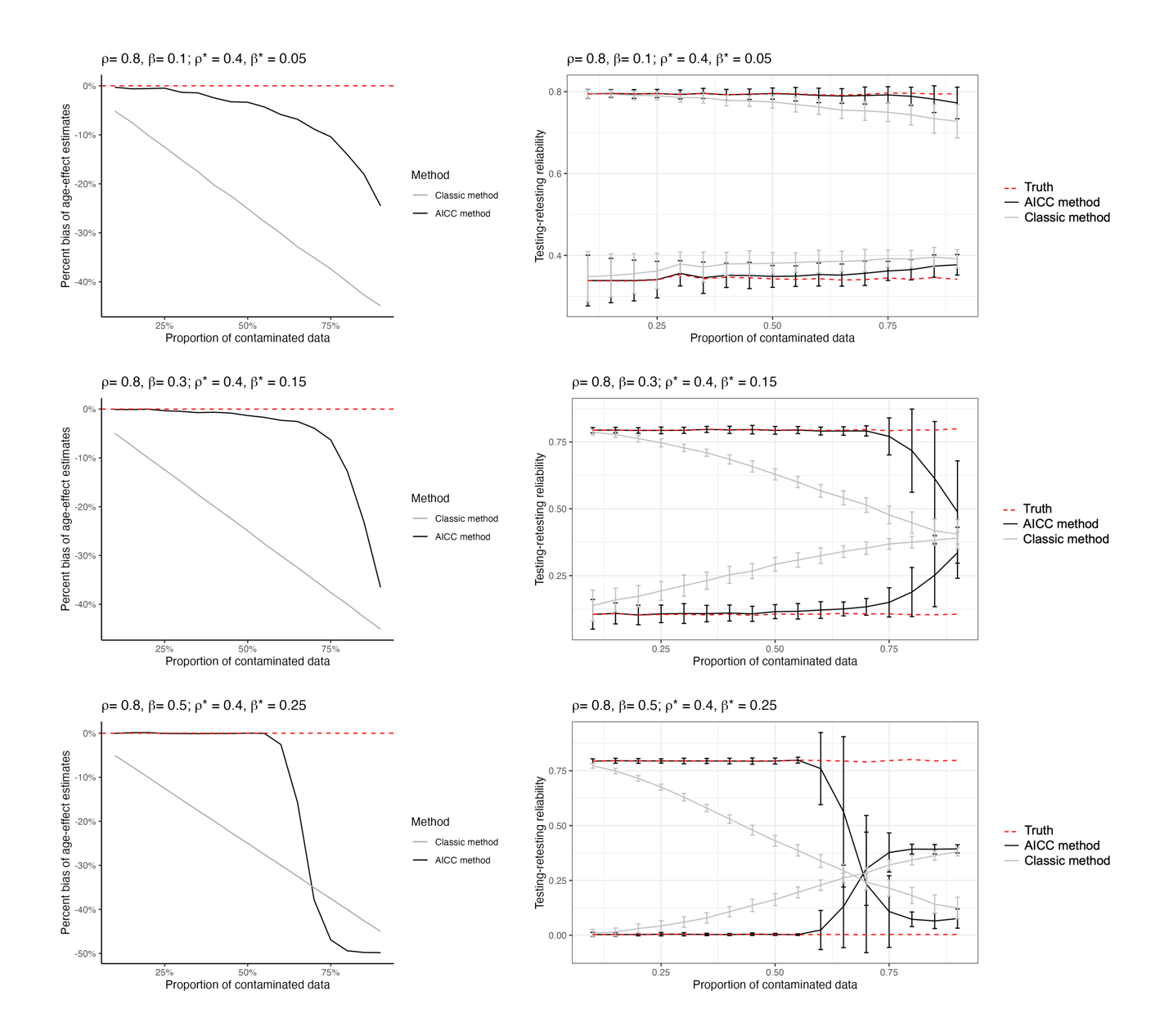

Figure S1. Simulation results. The left-side figures depict changes in the percent bias of estimated age effects across varying levels of contamination in the dataset and true population age effects. The percent bias of age effect estimation rises as the proportion of low-quality data increases within the dataset. Our proposed approach consistently outperforms the testing-retesting reliability measure obtained using classic mixed models when the proportion of contaminated data is below 70%. Moreover, the AICC estimated through our method demonstrates minimal bias when the proportion of low-quality data is less than 50%, regardless of β assumptions.

Reference

1.	Elyounssi S, Kunitoki K, Clauss JA, et al. Uncovering and mitigating bias in large, automated MRI analyses of brain development. *bioRxiv*. 2023/03/01/ 2023:2023.02.28.530498. doi:10.1101/2023.02.28.530498

2.	Fisher RA. On the "Probable Error" of a Coefficient of Correlation Deduced from a Small Sample. Journal article. *Metron*. 1921;1: 3-32doi:<http://hdl.handle.net/2440/15169>

3.	Shrout PE, Fleiss JL. Intraclass correlations: uses in assessing rater reliability. *Psychological Bulletin*. 1979/03// 1979;86(2):420-428. doi:10.1037//0033-2909.86.2.420
